# Heterogeneity among estimates of the core genome and pan-genome in different pneumococcal populations

**DOI:** 10.1101/133991

**Authors:** Andries J van Tonder, James E Bray, Keith A Jolley, Sigríður J Quirk, Gunnsteinn Haraldsson, Martin CJ Maiden, Stephen D Bentley, Ásgeir Haraldsson, Helga Erlendsdóttir, Karl G Kristinsson, Angela B Brueggemann

## Abstract

**Background:** Understanding the structure of a bacterial population is essential in order to understand bacterial evolution, or which genetic lineages cause disease, or the consequences of perturbations to the bacterial population. Estimating the core genome, the genes common to all or nearly all strains of a species, is an essential component of such analyses. The size and composition of the core genome varies by dataset, but our hypothesis was that variation between different collections of the same bacterial species should be minimal. To test this, the genome sequences of 3,121 pneumococci recovered from healthy individuals in Reykjavik (Iceland), Southampton (United Kingdom), Boston (USA) and Maela (Thailand) were analysed.

**Results:** The analyses revealed a ‘supercore’ genome (genes shared by all 3,121 pneumococci) of only 303 genes, although 461 additional core genes were shared by pneumococci from Reykjavik, Southampton and Boston. Overall, the size and composition of the core genomes and pan-genomes among pneumococci recovered in Reykjavik, Southampton and Boston were very similar, but pneumococci from Maela were distinctly different. Inspection of the pan-genome of Maela pneumococci revealed several >25 Kb sequence regions that were homologous to genomic regions found in other bacterial species.

**Conclusions:** Some subsets of the global pneumococcal population are highly heterogeneous and thus our hypothesis was rejected. This is an essential point of consideration before generalising the findings from a single dataset to the wider pneumococcal population.

## Background

Collectively, the complete set of genes possessed by members of a bacterial species is defined as the pan-genome [1], Understanding bacterial population structure requires knowledge of which genes in the pan-genome are found in all, or nearly all, strains of that species (core genes), and which are only found in some strains (accessory genes). In any study, investigators characterise a subset of the whole population; if one wishes to generalise the findings, then it must be determined whether or not the single dataset is likely to be representative of the whole population.

We developed a Bayesian decision model for estimating the bacterial core genome for datasets comprised of incomplete (draft) genome sequences generated via next-generation sequencing methodologies [2], In that study we included the estimation of core genomes for two different pneumococcal datasets, a diverse global historical dataset and a dataset of carriage genomes from Boston, Massachusetts, USA [3], More recently, two additional genome datasets of carriage pneumococci recovered from healthy children in Southampton, United Kingdom, and from young children and their mothers living in the Maela refugee camp on the Thailand-Myanmar border were published [4-5]. The genomes of pneumococci recovered from healthy young children recruited to our ongoing vaccine impact study in Iceland were also available, many of which have already been published [6-7], Thus four well-sampled pneumococcal genome datasets from four different geographical locations were available for this study.

The aim of our study was to test the hypothesis that the estimated core genome of any one dataset accurately represents the genes shared by pneumococci recovered in different geographical locations. To achieve this we analysed four datasets of carriage pneumococci and: i) estimated and compared the four individual core genomes; ii) identified and characterised the shared ‘supercore’ genome; and iii) assessed the genes that comprise the pan-genome of each dataset, with an emphasis on the Maela dataset.

## Results

### Estimated core genome comparisons

The study dataset was comprised of 3,121 genomes and each individual dataset represented a wide range of serotypes and clonal complexes (Table 1; Additional file 1). The number of dataset-specific core genes calculated for pneumococci recovered in Reykjavik (n = 1,059), Southampton (n = 1,052) and Boston (n =1,029) were nearly identical, but there were only 394 estimated core genes among Maela pneumococci (Table 2). For comparison, the number of core genes in a highly diverse global and historical dataset of 336 pneumococci recovered from both carriage and disease was estimated to be 851 genes using the same Bayesian model. The percentage of genomes in each dataset that possessed each estimated core gene ranged from ≥99.7% to ≥99.9%, which was consistent with the values calculated for other bacterial species datasets [2], The number of putative paralogues in any dataset was small and these were removed from further analyses.

**Table 1.**
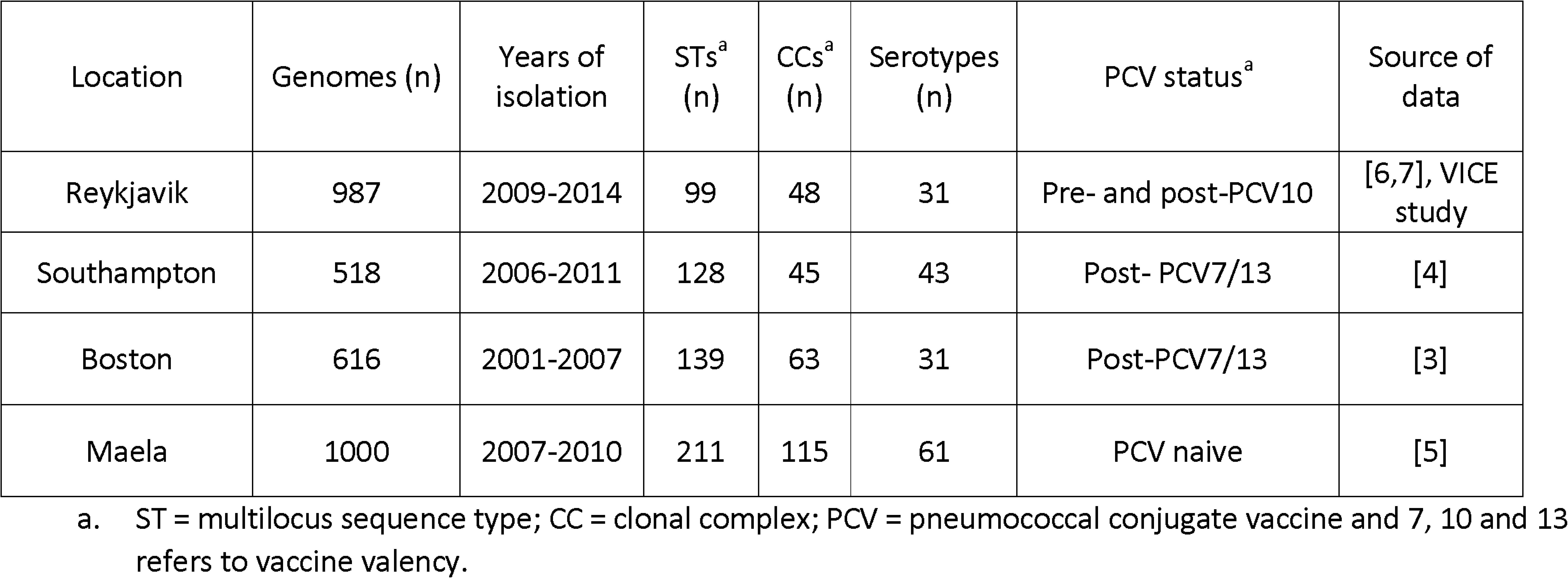
Summary of the pneumococcal genome datasets analysed in this study.

**Table 2.** Summary of the estimated core genome and pan-genome for each pneumococcal genome dataset.

| Location | % of genomes that possess each core gene | Putative paralogues (n) | Genes within estimated core genome (n) | Genes within pan-genome (n) |
| --- | --- | --- | --- | --- |
| Reykjavik | ≥99.9 | 8 | 1,059 | 7,301 |
| Southampton | ≥99.7 | 6 | 1,052 | 7,277 |
| Boston | ≥99.8 | 7 | 1,029 | 7,425 |
| Maela | ≥99.9 | 5 | 394 | 15,751 |

Despite the differences observed in the number of estimated core genes, the distribution of Clusters of Orthologous Groups (COG) functional categories among the core genes in each of the four datasets were similar (Figure 1; Additional file 2). In every case the largest proportion of estimated core genes were of unknown function (21.7-24.1%). Other major COG groups included genes associated with translation, ribosomal structure and biogenesis (11.9-15.7%), amino acid transport and metabolism (7.1-8.6%) and transcription (6.7-7.9%).

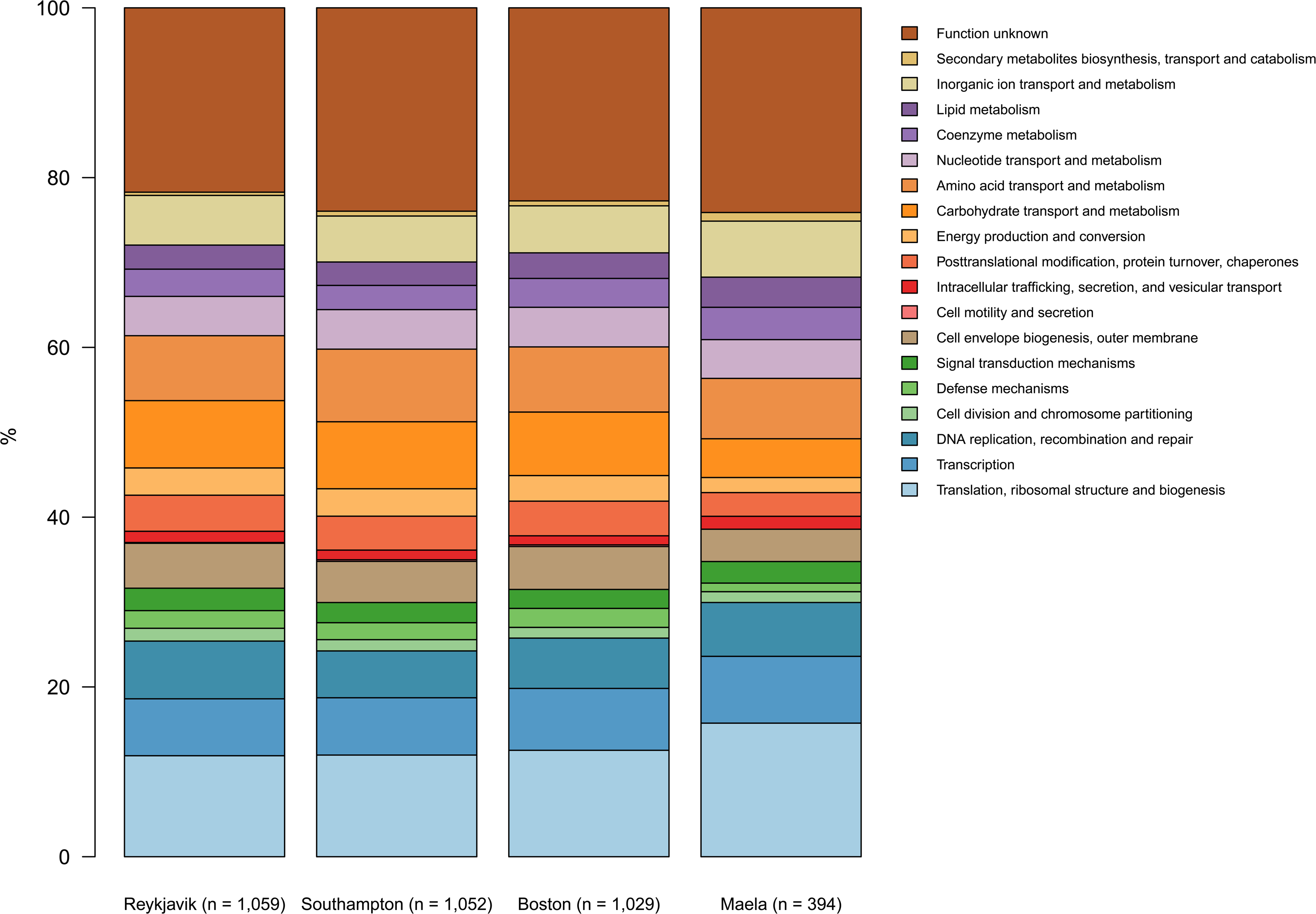

### The supercore genome and essential genes

There were 303 estimated core genes shared by all four pneumococcal datasets and we defined these as the supercore genome (Figure 2A). A further 461 genes were shared by the Reykjavik, Southampton and Boston pneumococci and thus there were 764 shared core genes in total between these three datasets (Additional file 3). Examination of the 461 genes common to the Reykjavik, Southampton and Boston datasets revealed that the distribution of COG functional categories broadly resembled that of the supercore genome (Figure 2B).

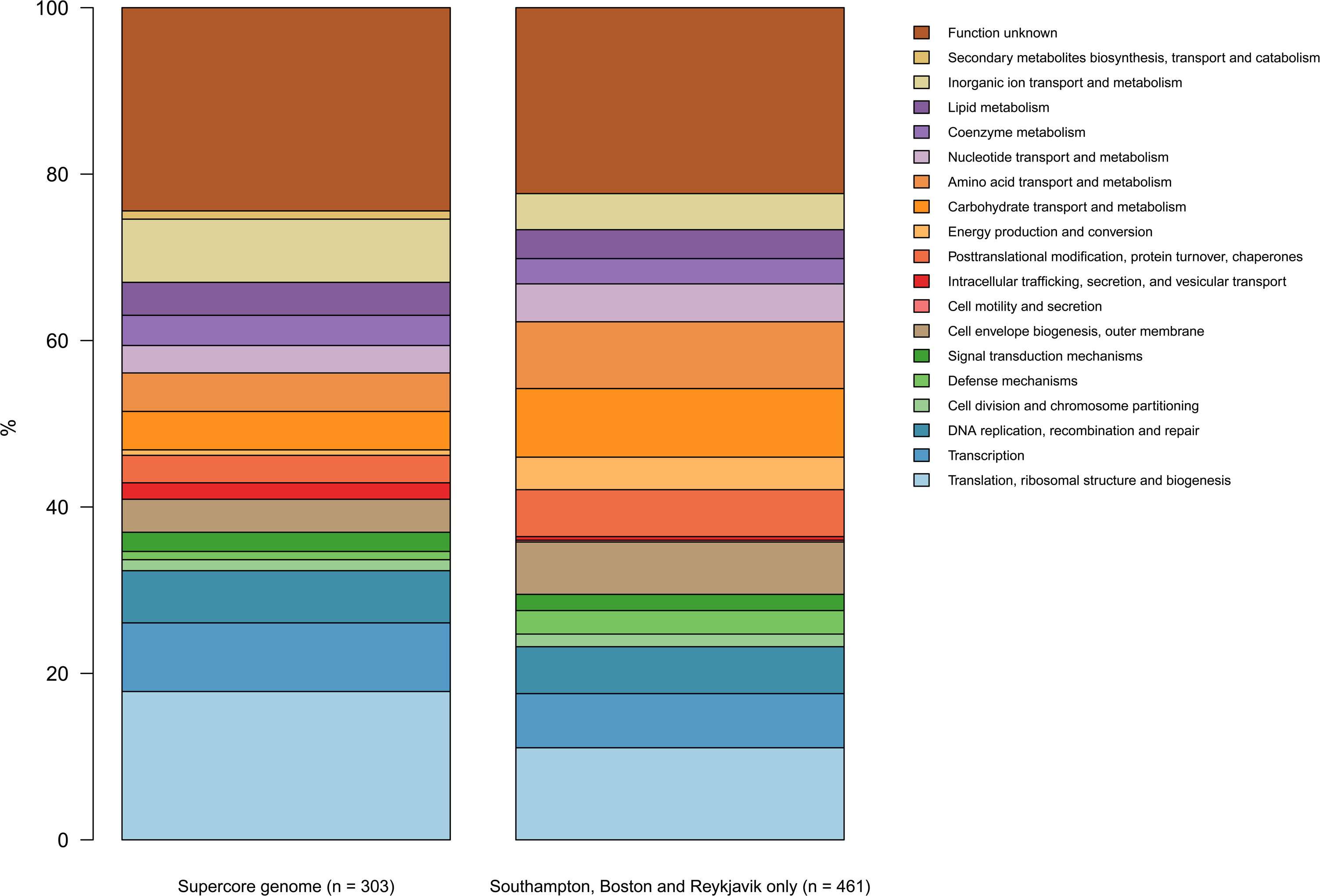

Earlier work by van Opijnen and colleagues predicted that 397 genes in an acapsular derivative of the TIGR4 pneumococcal genome were essential to fitness [8], but only 127 of these were amongst the supercore genes. The majority of these genes were involved in basic cell functions such as DNA replication, ribosomal proteins, RNA transcription and central carbon metabolism (Additional file 3).

### Supercore genome phylogeny

All 3,121 genomes were represented by a phylogenetic tree constructed using the 303 supercore gene sequences clustered with hierBAPS (Figure 3). The hierBAPS analysis revealed 19 monophyletic sequence clusters (SCs) that ranged in size from 36 to 263 genomes and were concordant with clonal complexes defined using MLST data. Pneumococci representing three clonal complexes were found in all four locations (CC^Predominant serotype(s)^): CC156/162^9V,19F^ (SC5); CC180^3^ (SC10); and CC448^NT^ (SC19). Pneumococci from other clonal complexes were identified in Reykjavik, Southampton and Boston, but not Maela: CC199^19A,15B/c^ (SCI); CC439^23F/B/A^ (SC2); CC395^6C^ (SC3); CC460^6A,10A,35F^ (SC6); CC62^11A^ and CC100^33F^ (SC8); CC433^22F^ (SC11); CC138/176^6B^ (SC13); and CC344^NT^ (SC18). All of these are widely-distributed genetic lineages [9].

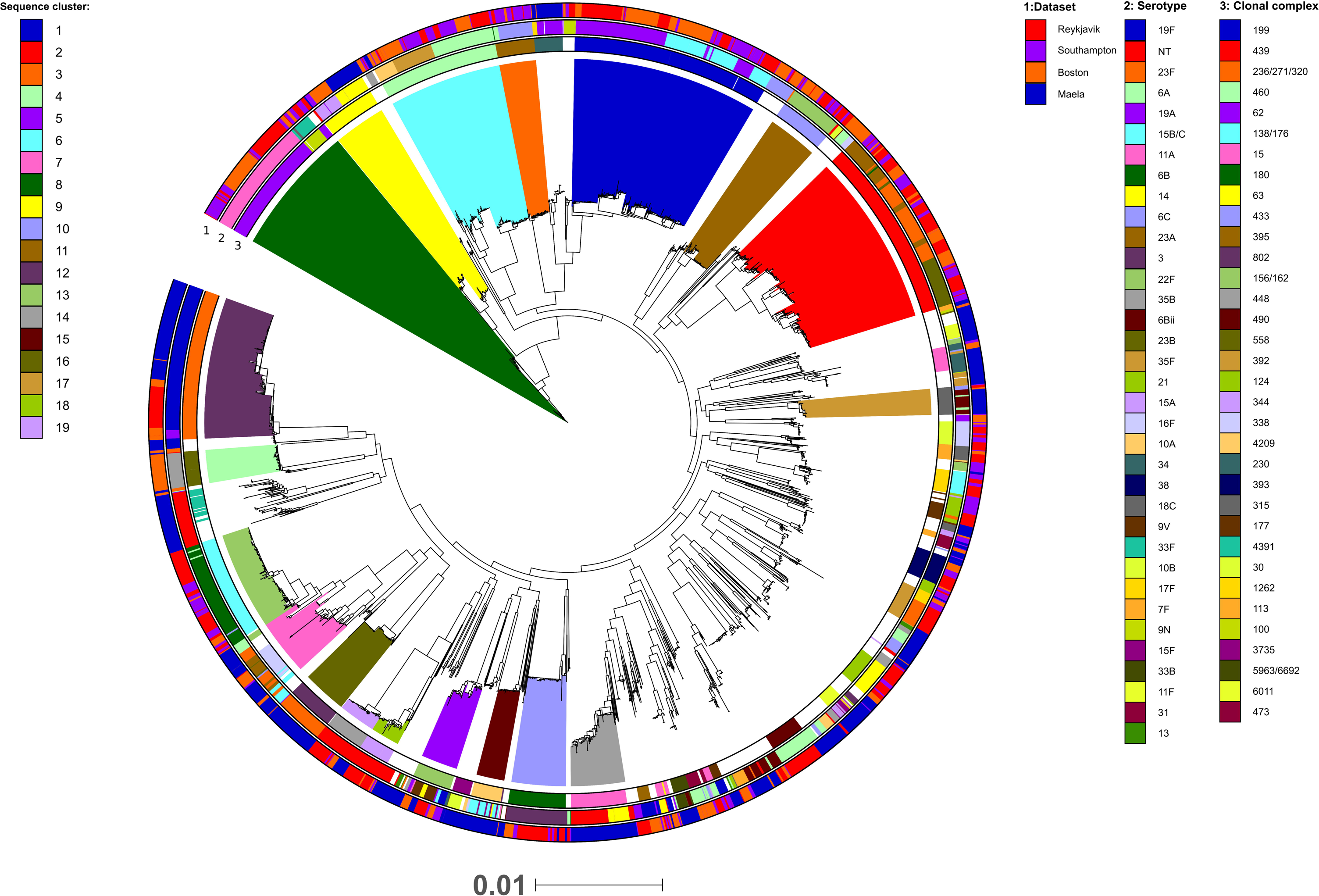

Pneumococci representing multidrug-resistant clonal complexes CC236/271/320^19F/A^ (SC12) and CC3 15^6Bii^ (SC17) were found in all locations apart from Southampton; CC63^14,15A^ (SC9) was found in all locations apart from Reykjavik. Conversely, pneumococci in CC15^14,NT^ (SC14), CC4209^15B/c^ (SC15) and CC802^23F^ (SC16) were only found in Maela, and pneumococci within CC558^35B^ (SC4) were only recovered in Boston.

SC7 was an unusual collection of 79 pneumococci from all four locations, although the majority were from Maela (n = 55) and Boston (n = 19). Six different clonal complexes were represented among pneumococci from Maela, three of which also represented the pneumococci from Boston (CC338^23F/A/B^, CC171^23F^ and CC138/176^15B/C,23A/F^). CC338^23F/A/B^ also represented the pneumococci in this cluster from Southampton (n = 3) and Reykjavik (n = 2).

The hierBAPS analysis also identified 1,177 genomes that were represented by polyphyletic sequence clusters (uncoloured clusters, Figure 3). More than half of the Maela genomes (55.1%) were part of these polyphyletic sequence clusters, in contrast to approximately 30% of the datasets from Reykjavik (30.1%), Southampton (30.5%) and Boston (27.8%).

### Pan-genome comparisons

The pan-genomes of the Reykjavik, Southampton and Boston pneumococci were very similar in size (7,277-7,425 genes), in contrast to the 15,751 genes in the Maela pneumococcal pan-genome (Table 2). The dataset-specific pan-genomes were calculated twice for each dataset, using ≥70% and ≥90% nucleotide sequence identity thresholds; however, there were minimal within-dataset differences in the total number of genes in the pan-genomes using either threshold (Figure 4A). The total number of genes in the pan-genomes of the Reykjavik, Southampton and Boston datasets plateaued at 6,000-7,000 genes, whilst the Maela pan-genome continued to increase. All four pan-genomes were open, i.e. the number of genes increased as more genomes were added to the analysis, which was a previously reported observation in pneumococcal pan-genome analyses [1].

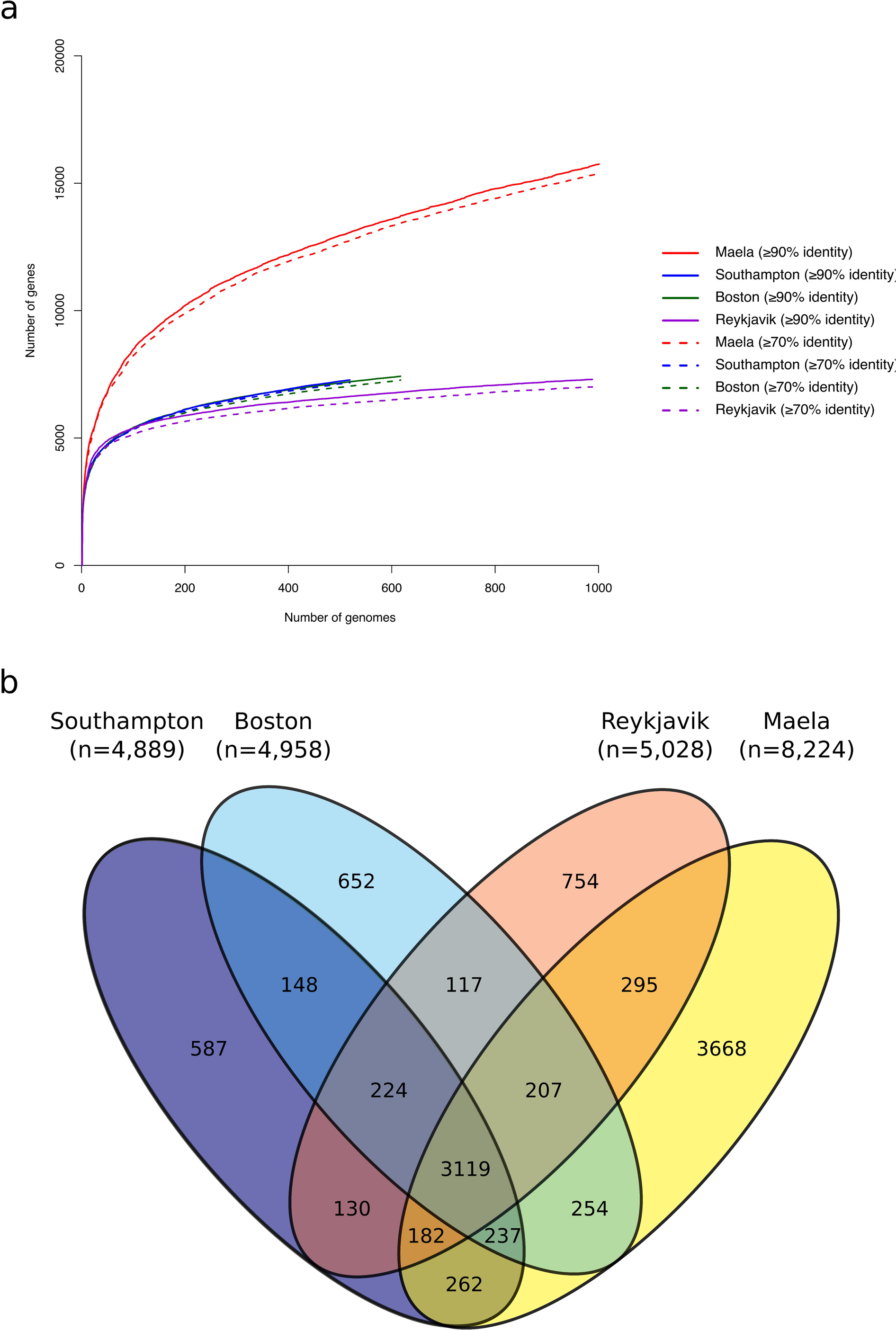

Overall, among the four datasets there were 37,754 genes in the combined pan-genome and these formed 10,836 gene clusters at a threshold of ≥70% nucleotide sequence identity. 3,119 of these gene clusters were identified among at least one pneumococcus from each of the four datasets (Figure 4B). The number of unique gene clusters in the Reykjavik (n = 754), Southampton (n = 587) and Boston (n = 652) datasets were broadly similar, as compared to 3,668 gene clusters unique to the Maela dataset. The function of nearly half of the unique gene clusters in any dataset was unknown (Additional files 4 and 5).

### Potential influence of nontypable pneumococci or non-pneumococcal *Streptococcus* spp. genomes

Nontypable pneumococci comprised 16.6% (n = 512) of the original Maela dataset, as compared to ≤6.6% of each of the three other datasets, and nontypable pneumococci are recognised as being a diverse group [10]. To test whether the inclusion of a large proportion of nontypable pneumococci were strongly influencing the findings, the nontypable pneumococci were excluded from the full Maela dataset, a random sample of 1,000 genomes was selected from the remaining genomes and the core genome and pan-genome were recalculated. Exclusion of the nontypable genomes had a minor effect on the size of the Maela pan-genome (decreased from 15,751 to 14,537 genes) and the estimated core genome (increased from 394 to 441 genes).

Another possible explanation for the observed differences was that the Maela dataset contained genomes from non-pneumococcal *Streptococcus* spp. To investigate this possibility, a phylogenetic tree was constructed based upon the 53 rMLST loci sequences extracted from the 3,121 pneumococcal genomes plus 1,000 genomes of 65 different non-pneumococcal *Streptococcus* spp. (Additional file 6) [11], All 3,121 pneumococcal genomes clustered together, separate from the non-pneumococcal *Streptococcus* spp. (Additional file 7); therefore, the observed differences among the Maela dataset were unlikely to be explained by the inclusion of non-pneumococcal genomes.

### Large unique gene regions in the Maela pan-genome

Although the Maela dataset was comprised of bona fide pneumococci, the large number of accessory genes unique to the Maela dataset suggested that genomic regions that were non-pneumococcal in origin might be influencing the results. Gene names and genomic positions for the 3,668 gene clusters unique to the Maela pan-genome were extracted and manually inspected to identify large (>25 Kb) genomic regions. 14 regions that ranged from 25.2 to 66.1 Kb were revealed, most of which were identified in multiple Maela genomes (Table 3). The nucleotide sequences of these 14 regions were extracted and used to query GenBank and the dataset of 1,000 non-pneumococcal *Streptococcus* spp. genomes to identify possible matches.

**Table 3.**
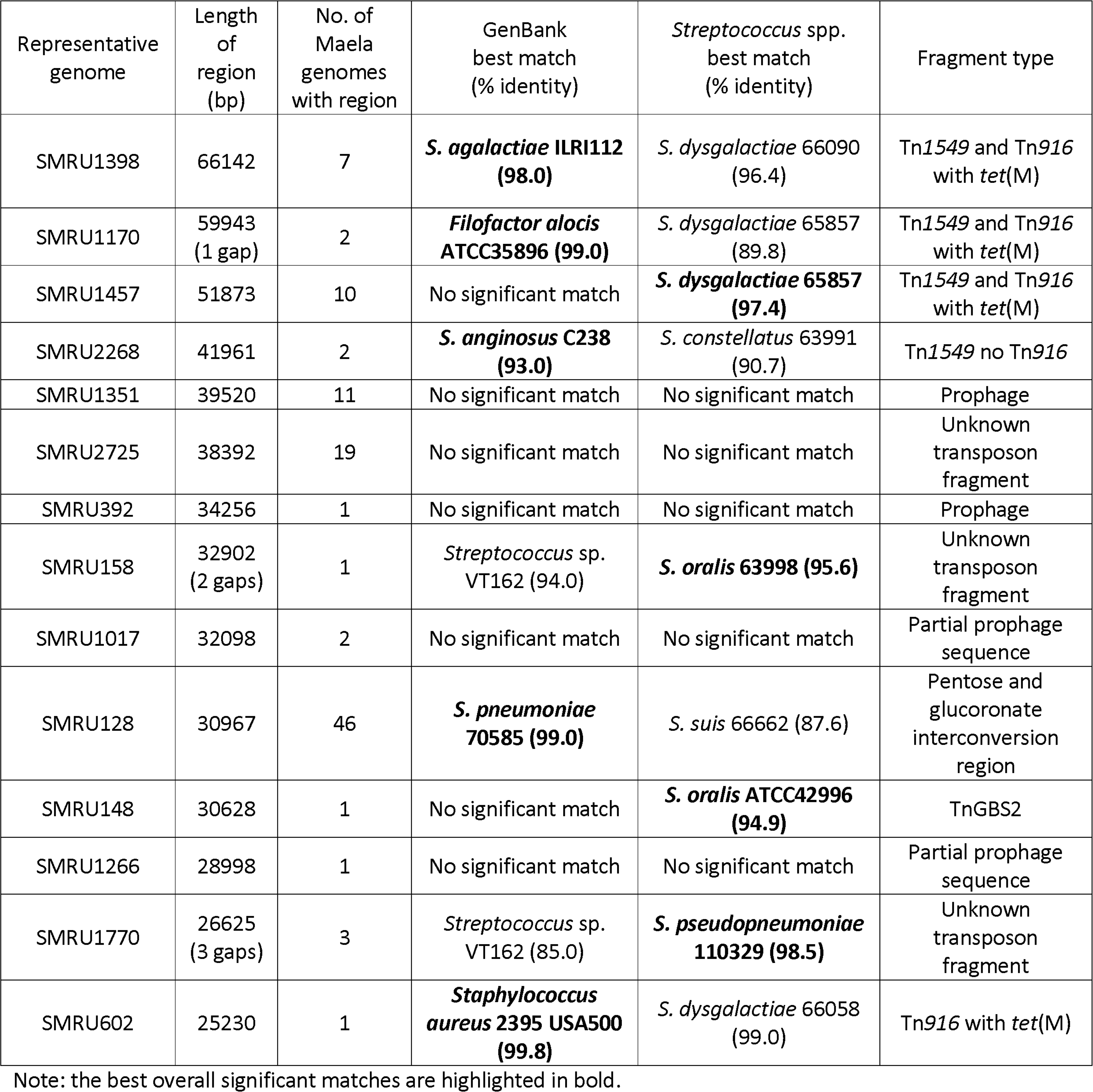
Large genomic regions that were unique to the Maela dataset.

| Representative genome | Length of region (bp) | No. of Maela genomes with region | GenBank best match (% identity) | *Streptococcus* spp. best match (% identity) | Fragment type |
| --- | --- | --- | --- | --- | --- |
| SMRU1398 | 66142 | 7 | ***S. agalactiae* ILRI112 (98.0)** | *S. dysgalactiae* 66090 (96.4) | Tn1549 and Tn916 with *tet*(M) |
| SMRU1170 | 59943 (1 gap) | 2 | ***Filofactor alocis* ATCC35896 (99.0)** | *S. dysgalactiae* 65857 (89.8) | Tn1549 and Tn916 with *tet*(M) |
| SMRU1457 | 51873 | 10 | No significant match | ***S. dysgalactiae* 65857 (97.4)** | Tn1549 and Tn916 with *tet*(M) |
| SMRU2268 | 41961 | 2 | ***S. anginosus* C238 (93.0)** | *S. constellatus* 63991 (90.7) | Tn1549 no Tn916 |
| SMRU1351 | 39520 | 11 | No significant match | No significant match | Prophage |
| SMRU2725 | 38392 | 19 | No significant match | No significant match | Unknown transposon fragment |
| SMRU392 | 34256 | 1 | No significant match | No significant match | Prophage |
| SMRU158 | 32902 (2 gaps) | 1 | *Streptococcus* sp. VT162 (94.0) | ***S. oralis* 63998 (95.6)** | Unknown transposon fragment |
| SMRU1017 | 32098 | 2 | No significant match | No significant match | Partial prophage sequence |
| SMRU128 | 30967 | 46 | ***S. pneumoniae* 70585 (99.0)** | *S. suis* 66662 (87.6) | Pentose and glucuronate interconversion region |
| SMRU148 | 30628 | 1 | No significant match | ***S. oralis* ATCC42996 (94.9)** | TnGBS2 |
| SMRU1266 | 28998 | 1 | No significant match | No significant match | Partial prophage sequence |
| SMRU1770 | 26625 (3 gaps) | 3 | *Streptococcus* sp. VT162 (85.0) | ***S. pseudopneumoniae* 110329 (98.5)** | Unknown transposon fragment |
| SMRU602 | 25230 | 1 | ***Staphylococcus aureus* 2395 USA500 (99.8)** | *S. dysgalactiae* 66058 (99.0) | Tn916 with *tet*(M) |

Note: the best overall significant matches are highlighted in bold.

Four different examples of Tn*l549*-like integrative and conjugative elements (ICEs), three of which included Tn*916* with *tet*(M), which mediates tetracycline resistance, were identified in 21 genomes (Table 3; Figure 5). The nearest matches to these variable Tn*l549*-like regions were predominately from non-pneumococcal *Streptococcus* spp., but a nearly identical match to the Tn*1549* region in one pneumococcal genome was from *Filofactor alocis* ATCC35896, a grampositive anaerobe implicated in periodontal disease [12]. Tn*916* was also found on its own in one Maela genome and it was identical to a Tn*916* in the *Staphylococcus aureus* 2395 USA500 genome. Tn*GBS2,* another type of ICE, was found in a single Maela genome. Tn*GBS2* uses a DDE transposase instead of a phage-like integrase for mobility and is found in oral *Streptococcus* spp. such as *S*. *mitis* and *S*. *oralis* [13].

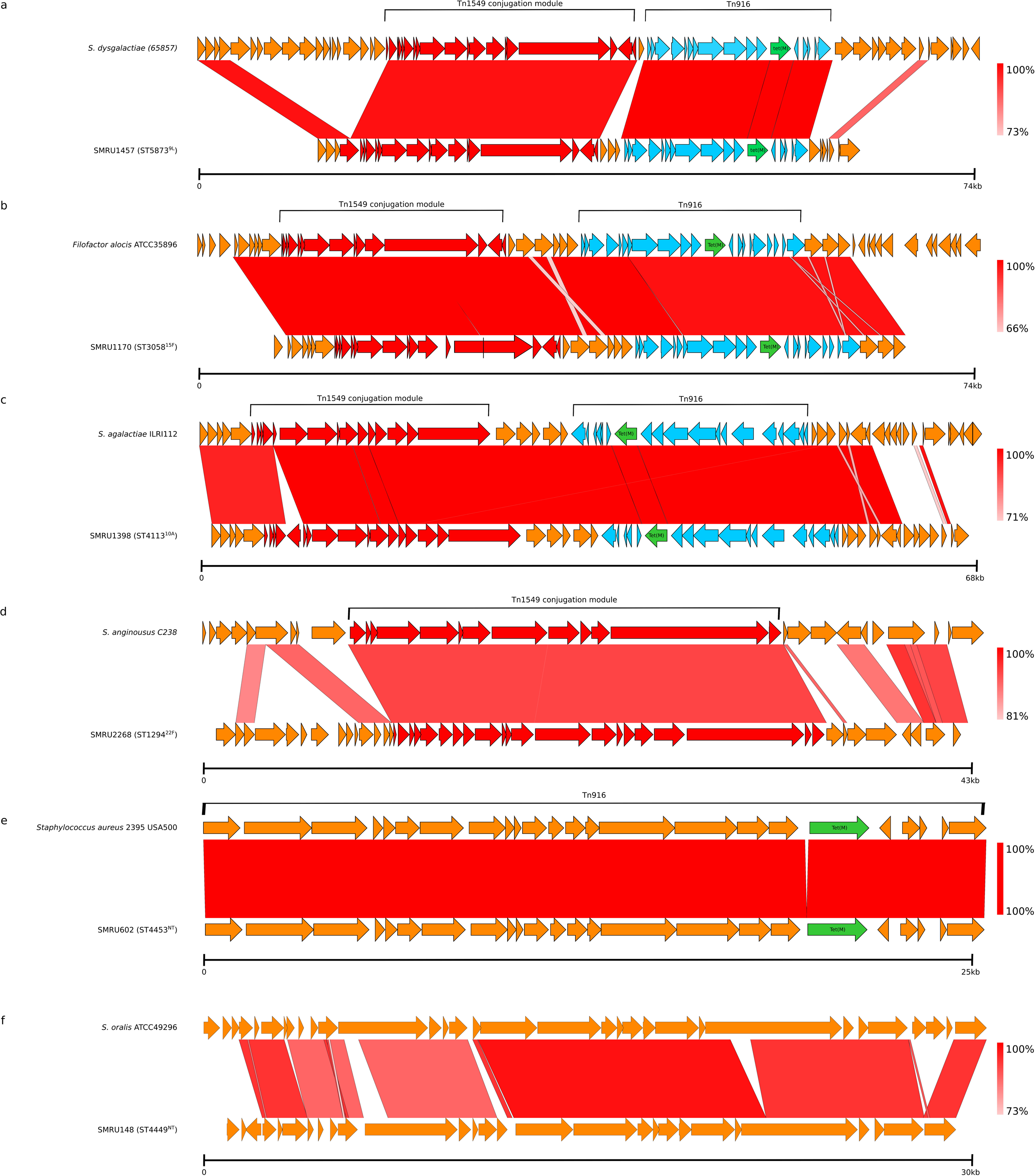

No significant matches were identified in either GenBank or the *Streptococcus* spp. genomes for four different prophage sequences (28.9-39.5 Kb) and an unknown transposon fragment. The four prophage sequences were compared to our database of pneumococcal prophage sequences assembled from another collection of diverse pneumococcal genomes. Two of the Maela sequences were of two different, putatively full length novel prophages that did not match to any previously reported prophages, but were related to one cluster E prophage from our recent study [14]. These were submitted to GenBank (accession numbers pending). The two other putative prophage sequences were of incomplete prophages with no close matches in our database.

46 genomes from multiple genetic lineages possessed a 31.0 Kb region that contained a number of genes involved in carbohydrate metabolism, including PTS lactose and ascorbate transporters, and genes that constituted the pentose and glucuronate interconversion pathway, which is an alternative to glycolysis [15]. A GenBank search revealed a nearly identical hit to S. *pneumoniae* 70585 (ST289^5^), a disease-causing pneumococcus from Bangladesh.

## Discussion

The relative ease with which bacterial genomes can be sequenced using next-generation sequencing technologies has resulted in a paradigm shift in our understanding of bacterial populations. MLST was developed nearly twenty years ago and it quickly became a powerful tool for defining bacterial lineages. The explosion of MLST data fundamentally informed our understanding of bacterial population structure, recombination, evolution, epidemiology, pathogenesis, and the consequences of perturbing bacterial populations. Genomics now provides the data with which one can address hypotheses with a much higher resolution than ever before. Genomics has not abrogated the relevance of MLST, but in fact genome sequence-based clustering (at least in the case of pneumococci) is highly concordant to clustering based on MLST data. This is helpful, since the MLST nomenclature is even more valuable with the overlay of genome-wide information, and as a result it is becoming clear that the diversity within some bacterial populations may be even more nuanced than previously appreciated.

Our study has clearly shown that geographically-distinct datasets of carried pneumococci from Reykjavik, Southampton and Boston were similar in terms of the size and composition of their estimated core genomes and overall pan-genomes. In contrast, the dataset from Maela was distinctive in terms of the large size of its pan-genome and small estimated core genome, as well as the overall diversity of its CC and serotype distributions. Maela is a refugee camp of only ∼50,000 inhabitants and the movement of people in and out of the camp is restricted; therefore, our expectation was that there would be a similar bottleneck in the flow of pneumococci in and out of the camp, leading to a comparatively less diverse pneumococcal population. This was not observed: the diversity of pneumococci circulating in the Maela refugee camp was greater than that in any of the cities of Reykjavik, Southampton and Boston. There were twice as many serotypes and more than double the number of CCs among the carriage pneumococci in Maela as there were in Reykjavik, which was the least diverse of the four datasets based on those criteria.

There were approximately five times as many unique gene clusters in the Maela pan-genome as there were in the other datasets. The large regions that were identified were predominately from other *Streptococcus* spp., but in two cases the best matches were to non-streptococcal bacteria. It seems likely that there are other genomic regions of interest to be found in the list of ∼3,700 genes unique to the Thailand pan-genome, thus a more in-depth study of these regions and any other large regions identified in the pan-genomes of the other three datasets should be performed in future work. Altogether, these findings raise an important point: the global pneumococcal population is likely to be more heterogeneous than currently appreciated, particular among those pneumococci from geographical regions that have never or rarely been sampled to date. Moreover, our findings suggest that caution should be exercised when inferring broader biological conclusions about the pneumococcus based on a dataset from a narrow population sampling.

The majority of epidemiological studies that have been conducted in developed countries have shown that the pneumococcal serotypes and STs that circulate in carriage and disease are broadly similar across different populations [9], In contrast, recent epidemiological studies in places like Bolivia, Kenya, Malaysia and Nepal, which characterised pneumococci only by traditional MLST, demonstrated that whilst the most prevalent serotypes tend to be the same as those in the developed world, the diversity of STs/CCs was greater [16-19], Importantly, safe and effective pneumococcal conjugate vaccines (PCVs) are now used in many countries, but they significantly disrupt the pneumococcal population structure and this can have unpredictable consequences [3, 20-21], Therefore, characterising the pre-and post-PCV pneumococcal population structure is essential in order to identify the changes that occur. Whilst traditional MLST is still highly useful in that regard and will remain the genotyping method of choice in many parts of the world for some time, genomics provides a higher discriminatory level of resolution to such analyses and should be employed wherever possible.

## Conclusions

The availability of thousands of bacterial genomes means that meta-analyses of large datasets can now be undertaken in order to more precisely delineate bacterial population structure and the composition of the bacterial population in terms of the pan-genome, core and accessory genomes. More studies like this one will need to be carried out in order to better understand the heterogeneity of the global pneumococcal population in particular and bacterial populations more generally.

## Methods

### Pneumococcal carriage datasets selected for analyses

Icelandic pneumococci (n = 987) were recovered from the nasopharynx of healthy children 1-6 years old attending day care centres located in the greater capital area of Reykjavik, Kopavogur and Hafnarfjordur from 2009-2014 [6-7], Pneumococci from Boston, Massachusetts, USA (n = 616) were recovered from the nasopharynx of healthy children <7 years old who were attending primary care facilities in and around Boston from 2001-2007 [3], Pneumococci from the United Kingdom (n = 518) were isolated from the nasopharynx of children ≤4 years old attending the Southampton General Hospital outpatient department from 2006-2011 [4], Nasopharyngeal pneumococci (n = 3,085) from Maela, a refugee camp close to the border of Thailand and Myanmar, were collected from a cohort of 528 infants and 242 of their mothers from 2007-2010 as part of a longitudinal carriage study [5], All of the pneumococcal genomes were sequenced on the Illumina platform and assembled at the Sanger Institute.

Children in Reykjavik, Southampton and Boston were vaccinated with the 7-, 10- or 13-valent pneumococcal conjugate vaccine (PCV) at some point before and/or after the time pneumococci were collected in each of the original studies. PCV10 was introduced into Iceland in 2011; PCV7 was used in the UK from 2006-2009 and PCV13 thereafter; and PCV7 was introduced in the USA in 2000 and was replaced by PCV13 in 2010 [3-4, 22], No PCV was used in Thailand at the time pneumococci were collected in Maela.

1000 genomes from the original Maela dataset were randomly selected (using R) for inclusion in this study, to avoid bias due to the large size of the Maela dataset and to select a dataset similar in size to that of Reykjavik [23], Metadata for the Southampton, Boston and Maela genome datasets were manually extracted from the original publications. Complete lists of the pneumococcal genomes included in this study, with accession numbers and available metadata are listed in Additional file 1 and all assembled genomes are available for download from PubMLST [9],

### Sequence types (STs), clonal complexes (CCs) and serotypes

Multilocus sequence type (MLST) data were auto-extracted from each genome using BIGSdb and STs were clustered into CCs using Phyloviz [24-25], seqSerotyper.R was used to assign serotypes based upon the nucleotide sequence of the capsular locus [26],

### Core genome analyses

Prokka was used to predict and annotate the coding sequences (CDS), hereafter referred to as ‘genes’ for simplicity, in each genome [27], Gene annotation was based upon a bespoke pneumococcal sequence database compiled for this study, which used the gene annotation data from all available pneumococcal genomes in GenBank [28], The resulting annotation files in gff format were input into Roary and clustered using sequence identity thresholds of ≥70% and ≥90% (the lower threshold to account for large nucleotide differences between the same gene in a population, e.g. nucleotide similarity of *pbp2x* may differ by ≥25% between penicillin-susceptible and -resistant pneumococci) [29], Core genomes were calculated for each dataset using our Bayesian method [2], Putative paralogues were removed and the resulting core genes were extracted and aligned using MAFFT [30], A Venn diagram was created to depict the number of core genes in each of the four datasets.

Four dataset-specific sets of core gene sequences were created by extracting one sequence for every core gene in each dataset. The four sets of core genes were then compared and clustered in cd-hit using a similarity threshold of ≥90% and the ‘supercore’ genome (core genes that were present in every dataset) was determined [31], COG functional groups were assigned to each gene using eggNOG [32],

Sequence alignments for the supercore genes were concatenated to create a supercore genome alignment that was used to create a phylogenetic tree using FastTreeMP [33], The tree was reconstructed to account for recombination using ClonalFrameML [34], Sequence clusters were delineated using hierBAPS and depicted on the final phylogenetic tree using iTOL [35-36],

### Pneumococcal essential genes

A recently published study used Tn-seq to identify pneumococcal genes likely to be essential for survival [8], The relevant amino acid sequences for these 397 putatively essential genes were extracted from the TIGR4 genome and cd-hit was used to compare the amino acid sequences of the essential genes and the supercore genes at a sequence identity threshold of ≥70%.

### Pan-genome analyses

The nucleotide gene sequences for each of the dataset-specific pan-genomes were clustered in cd-hit using a similarity threshold of ≥70% and an alignment threshold of ≥90%. The numbers of shared and unique genes in the pan-genome of each dataset were represented by a Venn diagram constructed using a custom script.

### Genome sequence quality and sampling strategy of the Maela dataset

The strikingly different results for the Maela pneumococci (see Results) prompted further analyses. All of the genomes for the Maela dataset were downloaded from the ENA and assembled using Velvet. Genome sequence assemblies were assessed for total genome length and number of contigs, and ribosomal MLST (rMLST) loci were tagged to assign the bacterial species [11], Among the Maela genome assemblies, 80 of the 3,085 genomes from the original dataset failed the initial quality control and were discarded. A subsequent examination of the sequence assembly metrics for the 1,000 randomly-selected genomes included in the current study revealed that one genome was poorly assembled (∼1,600 contigs). To test whether this genome significantly skewed the Roary pan-genome analyses, it was replaced by another genome of the same ST, serotype and rMLST type and the analyses were repeated. The results were unaffected: the core-genome increased by 4 genes and the pan-genome decreased by 64 genes (data not shown).

We compared the distribution of contigs assembled for all genome sequences and noted that there were differences between datasets: the Reykjavik genomes were assembled with the fewest number of contigs (range 9-219, mode = 33 contigs); Southampton (range 52-248, mode = 87 contigs) and Boston (range 50-246, mode = 88 contigs) were very similar; and the genomes in the Maela dataset were comprised from the largest number of contigs (range 83-1687, mode = 202 contigs), although the distributions of contigs overlapped (Additional file 8).

To check whether the random sampling strategy had somehow biased the Maela dataset to be more diverse, a further 1,000 genomes from the remaining ∼2000 genomes were sampled in the same manner. The overall number of STs and serotypes for this sample were nearly identical to the original sample dataset (data not shown) suggesting that the observed epidemiological diversity was similar to that of the other Maela genomes not included here.

Additionally, 1,000 genomes from 65 non-pneumococcal *Streptococcus* species were selected for comparative analyses to ensure that only pneumococci were included in this study (Additional file 6). BIGSdb was used to extract the rMLST gene sequences from the 1,000 *Streptococcus* genomes and 3,121 pneumococcal genomes: these sequences were aligned, concatenated and used to construct a phylogenetic tree [24,30, 33], ClonalFrameML was used to reconstruct the tree and annotation was performed with iTOL.

The 3,668 gene clusters unique to the Maela pan-genome were manually inspected using the gene identifier numbers assigned by Prokka and the gene frequency information provided by the Roary output. The nucleotide sequence for each unique region was extracted using Artemis and both GenBank and the set of 1,000 non-pneumococcal genomes were queried to find homologous regions of sequence [11,37], Putative transposons were annotated using the *CONJscan* module [38], Homologous regions were compared using diagrams created with EasyFig [39],

## Declarations

## Acknowledgements

Not applicable.

## Availability of data and materials

The assembled genome sequences and corresponding metadata are available from the PubMLST website (https://pubmlst.org/spneumoniae/) and raw genome sequence data are via the NCBI Sequence Read Archive (see Additional file 1 for accession numbers).

## Author’s contributions

Conceived and designed the study: AJvT and ABB. Collected and processed the Icelandic pneumococci: SJQ, GH, AH, HE, KGK. Extracted DNA from the Icelandic pneumococci: AJvT. Sequenced and assembled the Icelandic pneumococcal genomes: SDB. Assembled bacterial genomes in the rMLST databases: JEB and KAJ. Managed and/or curated the rMLST and/or PubMLST databases: JEB, KAJ, MOM, ABB. Performed the analyses: AJvT and ABB. Wrote the manuscript: AJvT and ABB. All authors read and approved the final manuscript.

## Consent for publication

Not applicable.

## Ethics approval and consent to participate

Not applicable.

## Additional files

Additional file 1: Datasets analysed in this study.

Additional file 2: Estimated core genes in each dataset.

Additional file 3: Supercore genes and the additional core genes found only in three locations.

Additional file 4: COG functional categories for the set of genes unique to the pan-genome of each dataset.

Additional file 5: Unique genes in the pan-genome of each dataset.

Additional file 6: Details of non-pneumococcal streptococci analysed in this study.

Additional file 7: Phylogenetic analysis of Streptococcus spp.

Additional file 8: Genome assembly contigs for each set of genomes.

